# ATP drives efficient terpene biosynthesis in marine thraustochytrids

**DOI:** 10.1101/2020.11.20.391870

**Authors:** Aiqing Zhang, Kaya Mernitz, Chao Wu, Wei Xiong, Yaodong He, Guangyi Wang, Xin Wang

## Abstract

Understanding carbon flux-controlling mechanisms in a tangled metabolic network is an essential question of cell metabolism. Secondary metabolism, such as terpene biosynthesis, has evolved with low carbon flux due to inherent pathway constraints. Thraustochytrids are a group of heterotrophic marine unicellular protists, and can accumulate terpenoids under the high salt condition in their natural environment. However, the mechanism behind the terpene accumulation is not well understood. Here we show that terpene biosynthesis in *Thraustochytrium* sp. ATCC 26185 is constrained by local thermodynamics in the mevalonate pathway. Thermodynamic analysis reveals the metabolite limitation in the nondecarboxylative Claisen condensation of acetyl-CoA to acetoacetyl-CoA step catalyzed by the acetyl-CoA acetyltransferase (ACAT). Through a sodium elicited mechanism, higher respiration leads to increased ATP investment into the mevalonate pathway, providing a strong thermodynamic driving force for enhanced terpene biosynthesis. The proteomic analysis further indicates that the increased ATP demands are fulfilled by shifting energy generation from carbohydrate to lipid metabolism. This study demonstrates a unique strategy in nature using ATP to drive a low-flux metabolic pathway, providing an alternative solution for efficient terpene metabolic engineering.

**IMPORTANCE:** Terpenoids are a large class of lipid molecules with important biological functions, and diverse industrial and medicinal applications. Metabolic engineering for terpene production has been hindered by the low flux distribution to its biosynthesis pathway. In practice, a high substrate load is generally required to reach high product titers. Here we show that the mevalonate-derived terpene biosynthesis is constrained by local pathway thermodynamics, which can only be partially relieved by increasing substrate levels. Through comparative proteomic and biochemical analyses, we discovered a unique mechanism for high terpene accumulation in marine protists thraustochytrids. Through a sodium induced mechanism, thraustochytrids shift their energy metabolism from carbohydrate to lipid metabolism for enhanced ATP production, providing a strong thermodynamic driving force for efficient terpene biosynthesis. This study reveals an important mechanism in eukaryotes to overcome the thermodynamic constraint in low-flux pathways by increased ATP consumption. Engineering energy metabolism thus provides an important alternative to relieve flux constraints in low-flux and energy-consuming pathways.

## INTRODUCTION

Terpenoids are a large class of secondary metabolites produced by plants and microbes (1). Two C_5_ terpene precursors, isopentenyl diphosphate (IPP) and dimethylallyl diphosphate (DMAPP), can be condensed into diverse classes of terpene molecules, including monoterpenes (C_10_), diterpenes (C_20_), triterpenes (C_30_), and so forth (2). Many terpenoids participate in a variety of biological processes, including cell wall biosynthesis, membrane function, electron transport, and conversion of light energy into chemical energy (3). The triterpene squalene (C_30_H_50_) is an important intermediate during cholesterol biosynthesis in eukaryotic cells (4). Squalene has shown great potential for medical applications, including suppressing colon carcinogenesis (5), improving immune system health (6), and enhancing drug and vaccine delivery as parenteral emulsions (7).

The livers of deep sea sharks contain 15-69 % by weight of squalene (8), and are one of the main commercial squalene sources. Metabolic engineering of microbes and plants thus serves as a valuable alternative to supply sustainable squalene to the market. However, engineering high terpene yield is challenging due to inherent flux constraints within the terpene biosynthesis pathway. In eukaryotes, the mevalonate pathway is mainly responsible for generating the terpene precursors IPP and DMAPP (9, 10). Three acetyl-CoA molecules are sequentially condensed into one IPP in six steps, consuming three ATP and two NADPH in total. IPP can then be converted to DMAPP by IPP isomerase (11). To further generate squalene, IPP and DMAPP are fused to form farnesyl diphosphate (FPP) by FPP synthase, followed by a head-to-head FPP addition into presqualene diphosphate (PSPP), and a reductive rearrangement to convert PSPP to squalene catalyzed by squalene synthase (12).

Two key factors govern the flux distribution into metabolic pathways, *i.e.* thermodynamics and enzyme kinetics. Due to the low carbon flux distribution into terpene biosynthesis, high substrate load is generally necessary to provide a favorable thermodynamic driving force for high terpene titers. Pathway yield can be further enhanced through kinetic improvement where rate-limiting enzymes are targeted for optimization of their expression levels. HMG-CoA reductase (HMGR) is widely recognized as the key enzyme in the mevalonate pathway to improve terpene flux through gene overexpression (13). Diverting carbon flux from competing metabolic processes into terpene biosynthesis can also effectively increase terpene flux (14). For intermediary terpene metabolites such as squalene, the loss of the product yield can be attributed to squalene consumption by downstream metabolism (15). Strategies such as product sequestering thus are often implemented to improve terpene yield. In plants, sequestering squalene into a lipid droplet has shown improved squalene yield (16).

Understanding the carbon flux controlling mechanism in an organism with native high terpene flux could provide valuable insight for terpene engineering. The marine protists thraustochytrids can accumulate high levels of terpenes such as squalene and carotenoids (17-19). The strain *Aurantiochytrium* sp. 18W-13a accumulated squalene up to 17 % of cell dry weight under optimized fermentation conditions (20). Several other thraustochytrids strains, including *A*. sp. TWZ-97 (21), *Schizochytrium mangrovei* PQ6 (22) and *S. limacinum* SR21 (23), also showed great potential for squalene production. In thraustochytrids, sodium was proposed to contribute largely to the osmotic adjustment but not cell physiology (24). In this study, we found that sodium elicits increased respiration rates in *Thraustochytrium* sp. ATCC26185 (hereafter *Thraustochytrium* sp.), leading to increased ATP investment in the mevalonate pathway and high squalene yield. The discovery of ATP directly driving terpene biosynthesis provides important insight for future terpene engineering efforts.

## RESULTS

### NaCl stimulates squalene accumulation during cell growth

Thraustochytrids require sodium for cell growth in the marine environment. To show how sodium affects cell growth and lipid metabolism in *Thraustochytrium* sp., we grew cells with (5 g/L NaCl) or without supplementing NaCl to the medium. Compared to cells grown in the sodium supplemented medium (NaCl-5), cells without sodium (NaCl-0) showed a small lag phase during the growth (Fig. 1A and 1B). However, the final biomass accumulation was similar under both conditions, reaching the maximum of ~ 12 OD after 72 hours of growth. On the other hand, the squalene accumulation was significantly higher with the sodium supplementation. Under both sodium conditions, the squalene titer in *Thraustochytrium* sp. reached the highest level in the beginning of stationary phase. The highest squalene titer was 123.6 mg/L in the NaCl-5 group (Fig. 1B), a 2-fold increase (*P* < 0.01) compared to the titer in the NaCl-0 group (Fig. 1A and 1B). The specific squalene productivity was highest (*P* < 0.01) after 60 hours of growth, reaching 4 and 2.1 mg/L/day/OD in the NaCl-5 and NaCl-0 groups, respectively (Fig. 1A and 1B).

**FIG 1.**
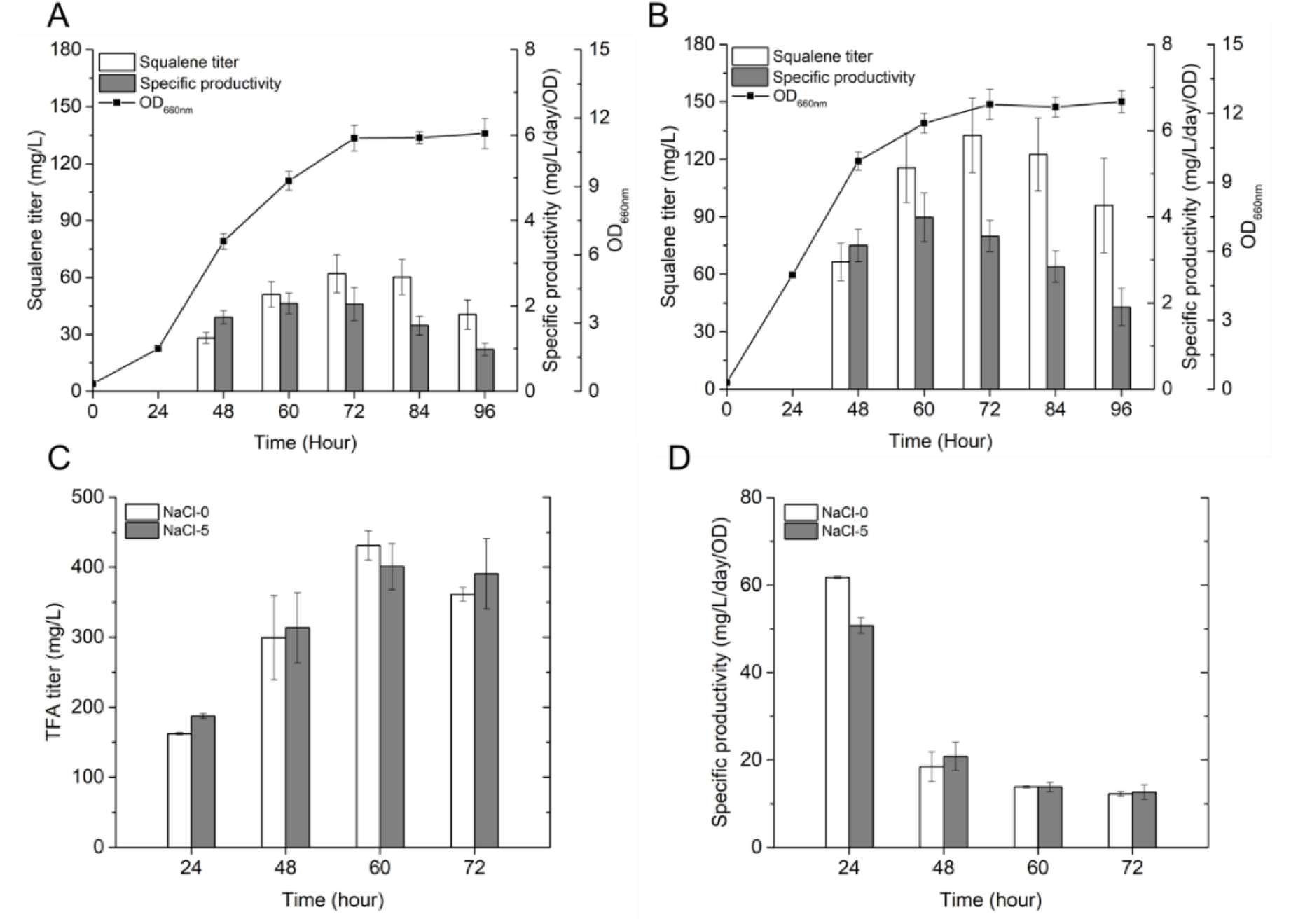
Growth and lipid production of *Thraustochytrium* sp. under different NaCl conditions. Squalene production and cell growth of cells cultivated (A) without NaCl (NaCl-0) and (B) with 5 g/L NaCl (NaCl-5). TFA titer (C) and specific productivity (D) of cells grown with or without NaCl.

Fatty acid biosynthesis consumes the same precursor acetyl-CoA as the mevalonate-derived squalene biosynthesis. If the increased squalene biosynthesis is due to higher acetyl-CoA levels under the salt condition, we could expect to see an increase in the total fatty acids (TFA). To understand how sodium affects fatty acid production, we measured the TFA contents in the NaCl-0 and NaCl-5 cells. The TFA analysis revealed the accumulation of a variety of fatty acids with different carbon lengths, including myristic acid (C14:0), pentadecylic acid (C15:0), palmitic acid (C16:0), margaric acid (C17:0), stearic acid (C18:0), and docosahexaenoic acid (C22:6) (Table S1). Although *Thraustochytrium* sp. cells showed a slight variation in their composition of C14, C15, and C16 fatty acids, both TFA titer and specific productivity were similar under two salt conditions (Fig. 1C and 1D and Table S1). The TFA titer had the highest level after 60 hours of cell growth, reaching 430.9 mg/L without NaCl and 401 mg/L with NaCl, respectively (Fig. 1C). The TFA specific productivity was highest at the 24-hour time point for both conditions, and decreased significantly when cells reached higher densities.

### NaCl induces a shift in energy metabolism that benefits ATP-consuming pathways

To understand how sodium affects cell metabolism in *Thraustochytrium* sp., we conducted comparative proteomics on cells grown under these two salt conditions. The cells with the highest specific squalene productivity (60 h) were used for the proteomic analysis. For each salt condition, an average of about 1,000 proteins were identified from the proteomics. Proline and glutamate are two commonly known compatible solutes for osmotic adjustment. In our proteomics, ornithine aminotransferase (OAT) and delta-1-pyrroline-5-carboxylate dehydrogenase (P5CDH), two key enzymes catalyzing the generation of proline and glutamate, showed significantly higher levels in cells grown with sodium (Fig 2), indicating their potential involvement in the osmotic adjustment in *Thraustochytrium* sp..

**FIG 2.**
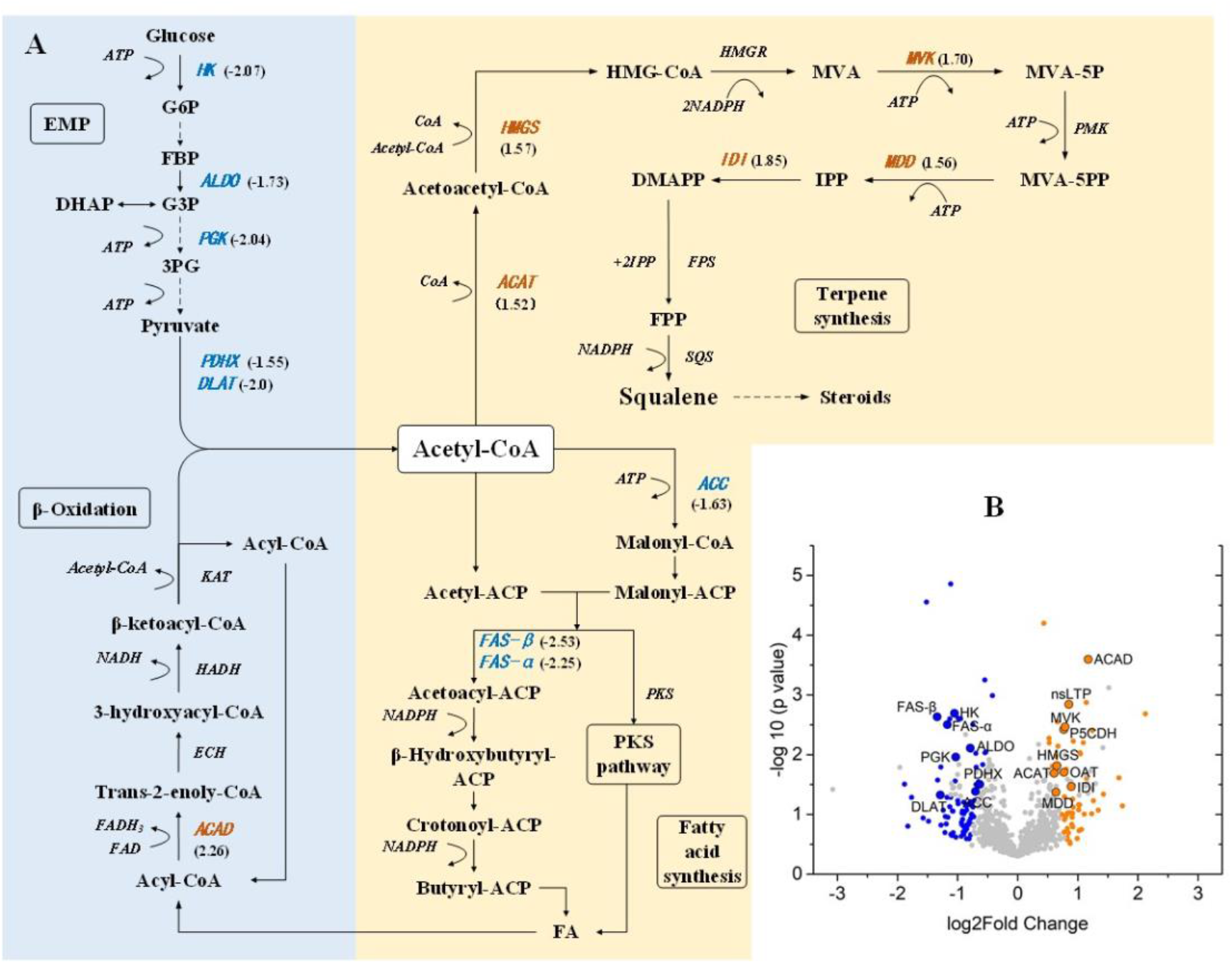
Sodium elicits an energy metabolism shift in *Thraustochytrium* sp.. **A.** Pathway diagram showing squalene synthesis from either glucose or lipid metabolism. Enzymes with statistically different levels in the comparative proteomics are shown in blue (decrease) and orange (increase) with the relative fold change in brackets. **Enzymes:** (HK: hexokinase, ALDO: fructose-bisphosphate aldolase, PGK: phosphoglycerate kinase, PK: pyruvate kinase, PDHX: pyruvate dehydrogenase protein X component, DLAT: dihydrolipoyllysine-residue acetyltransferase component of pyruvate dehydrogenase complex, ACAT: acetyl-CoA acetyltransferase, HMGS: 3-hydroxy-3-methylglutaryl-CoA synthase, HMGR: 3-hydroxy-3-methylglutaryl-CoA reductase, MVK: mevalonate kinase, PMK: phosphomevalonate kinase, MDD: diphosphomevalonate decarboxylase, ACAD: acyl-CoA dehydrogenases, ECH: enoyl-CoA hydratase, HADH: 3-hydroxyacyl-CoA dehydrogenase, KAT: 3-ketoacyl-CoA thiolase, IDI: isopentenyl-diphosphate isomerase, FPS: farnesyl diphosphate synthase, SQS: squalene synthase, FAS-α: fatty acid synthase subunit alpha, FAS-β: fatty acid synthase subunit beta). **Metabolites:** (G6P: D-glucose 6-phosphate, FBP: D-fructose 1,6-bisphosphate, G3P: D-glyceraldehyde 3-phosphate, 3PG: 3-phospho-D-glyceroyl phosphate, HMG-CoA: (S)-3-hydroxy-3-methylglutaryl-CoA, MVA: mevalonate, MVA-5P: mevalonate-5-phosphate, MVA-5PP: mevalonate-5-pyrophosphate, IPP: isopentenyl diphosphate, DMAPP: dimethylallyl diphosphate, FPP: farnesyl diphosphate, FA: fatty acid). **B**. Volcano plot of the whole proteome. Orange and blue dots represent proteins with significantly increased and decreased levels when sodium is supplemented in the growth medium. Grey dots represent proteins with similar protein levels under two growth conditions. **Enzymes and proteins**: (OAT: ornithine aminotransferase, P5CDH: delta-1-pyrroline-5-carboxylate dehydrogenase, nsLTP: nonspecific lipid transfer protein).

Under the salt condition, key enzymes in the Embden-Meyerhof-Parnas (EMP) pathway and fatty acid synthesis pathway showed significantly lower abundances compared to cells grown without sodium (Fig. 2A and 2B). Hexokinase (HK) and fructose-bisphosphate aldolase (ALDO) showed 2.1 and 1.7 -fold decreases in enzyme levels when cells grew under the salt condition. The protein abundance of phosphoglycerate kinase (PGK), the enzyme catalyzing one of the substrate-level phosphorylation steps in the EMP pathway, decreased 2-fold when cells were grown under the salt condition. In addition, two subunits of the pyruvate dehydrogenase complex, the enzyme responsible for the conversion of pyruvate to acetyl-CoA, had significantly lower abundances in cells grown under the salt condition, suggesting a lower flux toward acetyl-CoA generation through glucose catabolism. Interestingly, key enzymes for fatty acid biosynthesis also showed decreased levels in cells grown under the salt condition. Two pathways are responsible for fatty acid synthesis in thraustochytrids, *i.e.* the fatty acid synthase I (FAS I) system and the polyketide synthase-like (PKS-like) pathway (25). In cells grown under the salt condition, acetyl-CoA carboxylase (ACC), FAS I subunit alpha, and FAS I subunit beta showed 1.6, 2.3, and 2.5 -fold decreases in their protein abundances, respectively. The components of the PKS-like pathway did not show differences in their protein levels. The above observations suggest a decreased carbon flux from glucose catabolism and a lower activity toward fatty acid synthesis in *Thraustochytrium* sp. under the salt condition.

On the other hand, the acyl-CoA dehydrogenase (ACAD), a key enzyme in the β-oxidation pathway (26), showed a 2.2-fold enzyme increase in cells grown under the salt condition (Fig. 2A and 2B). The higher ACAD level suggests an enhanced ATP generation through the more efficient lipid metabolism when sodium is supplemented into the growth medium. A lipid-transfer protein also showed 1.8-fold protein increase under the sodium condition, suggesting its possible involvement in transferring fatty acids from cytosol to mitochondria for lipid degradation. These drastic changes in the enzymes associated with energy generation showed a clear shift from carbohydrate to lipid metabolism when the cells were grown under the salt condition.

The lipid metabolism through β-oxidation could also lead to higher acetyl-CoA levels, benefiting acetyl-CoA consuming processes such as the mevalonate derived terpene biosynthesis (27). Interestingly, the proteomic analysis showed significantly increased levels of MVA pathway enzymes in cells grown with sodium. Specifically, the enzyme abundances of acetyl-CoA acetyltransferase (ACAT), hydroxymethylglutaryl-CoA synthase (HMGS), mevalonate kinase (MVK), and mevalonate diphosphate decarboxylase (MDD), increased 1.5, 1.6, 1.7, and 1.6 -fold, respectively when cells were grown with sodium. These above results indicate a higher flux toward terpene biosynthesis under the salt condition. The energy metabolism shift from carbohydrate to lipid metabolism thus could potentially benefit terpene biosynthesis either through the increased acetyl-CoA input or ATP generation.

### *in silico* analysis reveals local thermodynamic constraint in squalene biosynthesis

The proteomic analysis suggests a shift in energy mechanism to facilitate the squalene synthesis in *Thraustochytrium* sp.. To understand why cells switch to lipid metabolism, we investigated the thermodynamic landscape of squalene biosynthesis using the pathway analysis tool PathParser (28). The PathParser is able to identify the step of a metabolic pathway whose flux is constrained by thermodynamics under physiological conditions. The pathway optimization process further seeks to maximize the driving force of the whole pathway by optimizing metabolite concentrations for all reactions within the defined concentration range (1 μM to 10 mM).

In the thermodynamic modeling, either glycolysis or β-oxidation of a short chain fatty acid hexanoyl-CoA was used as the upstream pathway for squalene synthesis. Interestingly, the optimized squalene synthesis pathway is constrained by the ACAT catalyzed reaction, regardless of the upstream catabolic processes (Fig. 3). ACAT catalyzes the nondecarboxylative Claisen condensation of two molecules of acetyl-CoA into acetoacetyl-CoA in the mevalonate pathway. The optimized ACAT reaction has a Δ*G*’ of −1.69 kJ/mol with glucose as the starting substrate, and a Δ*G*’ of near 0 kJ/mol when hexanoyl-CoA serves as the starting substrate (Table S2 and S3). In particular, the reaction is constrained by both the upper bound (10 mM) of acetyl-CoA concentration and the lower bound (1 μM) of acetoacetyl-CoA concentration as both metabolites are approaching their concentration limits under the optimized conditions (Table S4). Under physiological conditions, the metabolite concentration is seldom higher than 10 mM or lower than 1 μM, which indicates a thermodynamic limitation in the mevalonate-derived terpene biosynthesis pathway. In fact, if the concentration of coenzyme A increases to 5 mM instead of the defined 1 mM, the ACAT catalyzed reaction becomes infeasible with positive Δ*G*’ values for both glucose and hexanoyl-CoA derived squalene synthesis (Table S5, S6, S7). These analyses suggest that the ACAT catalyzed reaction operates near equilibrium under physiological conditions, and would require high enzyme kinetics (k_cat_/K_M_) in order to sustain a forward net flux (29).

**FIG 3.**
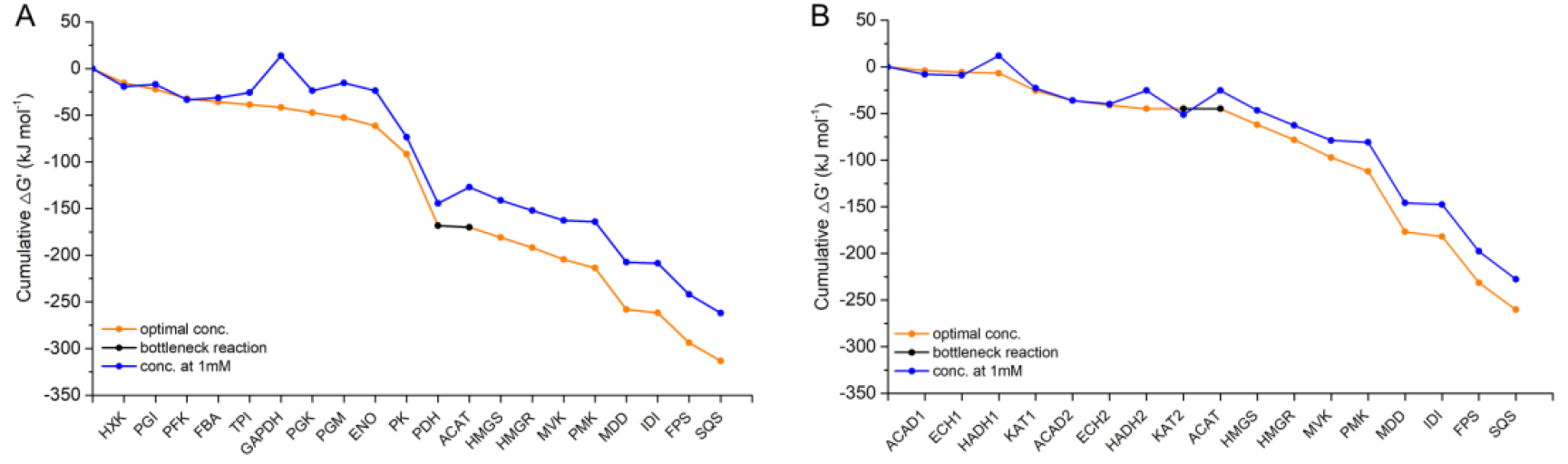
Thermodynamic landscape of squalene synthesis from carbohydrate and lipid metabolism. *in silico* analysis of squalene synthesis from glucose (A) and a short-chain fatty acid hyxanoyl-CoA (B). Blue line denotes standard Gibbs free energy at physiological 1 mM concentrations of metabolites. Orange line denotes reactions with optimized metabolite concentrations to provide maximal driving force for the whole pathway. The black line indicates the thermodynamic-constrained bottleneck reaction in the pathway.

Both glycolysis and β-oxidation can lead to the production of acetyl-CoA, which can be transported out of mitochondria for terpene synthesis in the cytosol. The thermodynamic analysis also indicates an inefficiency of driving the mevalonate pathway by increasing the acetyl-CoA concentration. In fact, the measured acetyl-CoA concentration in eukaryotic cells is in the micromolar range (30), hundreds of fold lower than the concentration used in the modeling. How can cell then benefit from switching to lipid metabolism to synthesize terpenoids? Besides acetyl-CoA, β-oxidation also generates reducing equivalents FADH_2_ and NADH that can directly participate in the electron transport chain for ATP production. On the other hand, the mevalonate pathway has three ATP-dependent steps catalyzed by MVK, phosphomevalonate kinase (PMK), and MDD. It is tempting to assume that efficient ATP production could provide a strong thermodynamic driving force for the mevalonate pathway, which can help alleviate the thermodynamic constraint in the ACAT catalyzed step.

### NaCl stimulates increased ATP production and drives efficient squalene synthesis

To verify whether ATP production can provide strong thermodynamic driving force for squalene biosynthesis, we first measured the ATP levels under the two salt conditions. Surprisingly, the ATP concentration remained at similar levels throughout cell growth (Fig. 4A), indicating a tight cellular regulation to achieve energy homeostasis. The measured ATP level represents only the steady state metabolite profile, and does not reflect its dynamic changes. We thus measured cellular respiration by looking at oxygen consumption rates. Under both growth conditions, the oxygen consumption rates decreased along with the cell growth. In addition, the oxygen consumption rates showed similar levels at the 24-hour time point between cells grown with and without sodium. However, the oxygen consumption rates were significantly higher in cells grown with sodium than those without sodium after 24 hours of cultivation. At all three time points (48 h, 60 h, and 72 h), the respiration rates of NaCl-5 cells were 1.7, 1.8, and 1.5 -fold higher, respectively in comparison with the NaCl-0 cells (Fig. 4B). This observation strongly supports our hypothesis that increased ATP generation leads to enhanced terpene production in *Thraustochytrium* sp..

**FIG 4.**
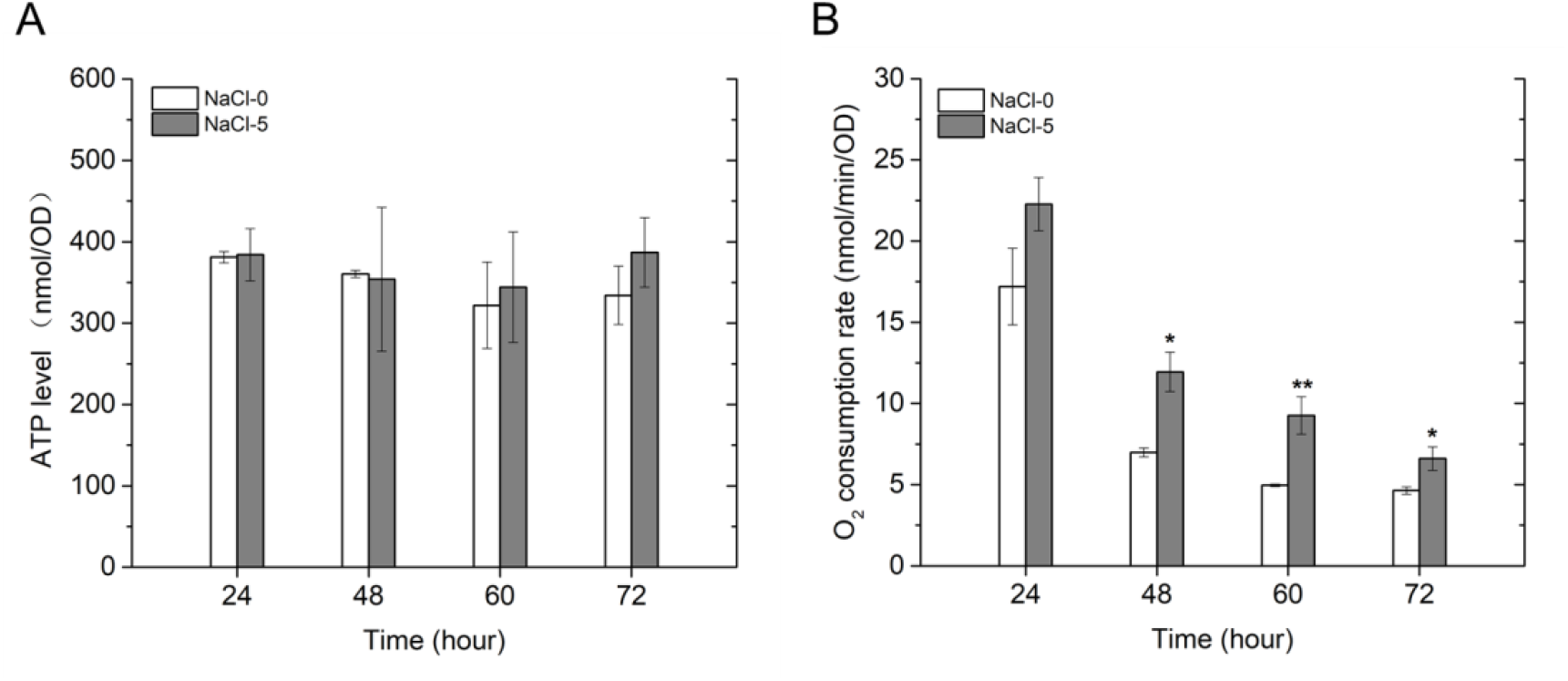
Energy status of *Thraustochytrium* sp. under different sodium conditions. ATP levels (A) and respiration rates (B) were measured along cell growth with and without NaCl supplemented in the medium.

## DISCUSSION

The yield of a target compound is determined by the carbon flux distributed to its biosynthesis pathway. Both enzyme kinetics and thermodynamics determine carbon flux distribution in metabolic pathways. The flux control coefficient (*C*) shows the relative contribution of a specific enzyme to the overall pathway flux. Kinetic bottlenecks can be alleviated through enzyme overexpression or substitution with enzyme homologs of higher activities (31). Contribution of thermodynamics to pathway flux is not widely appreciated. Not only will thermodynamics determine the feasibility of a reaction, it also constrains the pathway kinetics according to the flux-force relationship (29, 32). When a reaction approaches equilibrium, *i.e.* the Gibbs free energy change Δ*G*’ approaches 0, a large portion of the enzymes are used to catalyze the reverse reaction. A substantial increase in enzyme numbers is thus required to sustain the net flux. We show in this study that terpene synthesis is constrained by local pathway thermodynamics in the mevalonate pathway, which can be relieved by increased ATP production through a sodium elicited mechanism in *Thraustochytrium* sp.. Increased ATP consumption provides a strong thermodynamic driving force for the mevalonate pathway, leading to enhanced squalene biosynthesis in *Thraustochytrium* sp. (Fig. 1).

Pathway constraints by local thermodynamics are not uncommon in nature. For instance, many of the carbon fixation pathways in nature consume more ATP than what is necessary to make the overall reaction thermodynamically favorable. It was found that two types of reactions, *i.e.* carboxylation and carboxyl reduction, are energetically constrained, thus requiring activation from ATP hydrolysis directly or indirectly (33). The Milo group developed a quantitative framework for the analysis of pathway thermodynamics. This simple thermodynamically-derived metric, named the Max-min Driving Force (MDF) (29), can be used to compare flux capacity permitted by pathway thermodynamics under physiological conditions. In this study, we applied the MDF metric using the pathway analysis tool, PathParser (28), to identify steps constrained by metabolites levels under physiological conditions. In the mevalonate pathway, the ACAT catalyzed condensation of two acetyl-CoA into acetoacetyl-CoA is a thermodynamically-constrained bottleneck step (Fig. 3). Based on calculation using eQuilibrator (34), this reaction has an estimated Δ*G*°’ of +26 kJ/mol (pH = 7.5; ionic strength of 0.2 M), indicating a highly unfavorable reaction. Our thermodynamic analysis indicates that this reaction operates near equilibrium under defined physiological conditions, and easily becomes infeasible with the acetyl-CoA levels at the micromolar range found in eukaryotic cells (30) (Fig. 3 and Table S2-S7). The mevalonate pathway is an important metabolic process found in all three kingdoms of life (35). How do cells then solve this conundrum under physiological conditions? A recent study has shed some light into this problem. In the methanogenic archaeron *Methanothermococcus thermolithotrophicus,* a large enzyme complex forms to connect ACAT with HMGS, the enzyme catalyzing the addition of a third acetyl-CoA to acetoacetyl-CoA to form HMG-CoA in the mevalonate pathway (36). The enzyme complex forms a shared CoA-binding site, allowing substrate channeling between the endergonic nondecarboxylative Claisen condensation and the exergonic HMG-CoA-forming reaction, substantially improving the thermodynamics of the mevalonate pathway. Indeed, when these two reactions are coupled, the overall reaction to convert three acetyl-CoA to HMG-CoA has an estimated Δ*G*°’ of +4.7 kJ/mol calculated from eQuilibrator (34), which can be easily overcome by altering metabolite concentrations under physiological conditions. In the above study, genetic analysis also indicates the prevalence of this enzyme complex in archaea and many bacteria, but not in eukaryotes (36). Lone-standing ACAT and HMGS are widespread in eukaryotes and some bacteria, which might explain the low terpene flux observed in these organisms. On the other hand, archaea have high demands for terpenoids as membrane lipids, requiring an efficient mechanism to overcome the highly unfavorable acetyl-CoA condensation reaction.

In eukaryotic thraustochytrids, cells seem to overcome the thermodynamic limitation of the nondecarboxylative Claisen condensation through a different mechanism. The mevalonate pathway has three ATP-dependent reactions downstream of mevalonate formation. In our study, *Thraustochytrium* sp. has over 1.5-fold increases of respiration rates across the growth period when cells are cultured under sodium salt conditions (Fig. 4B). Two ATP-dependent enzymes MVK and MDD in the mevalonate pathway also have over 1.5-fold increases in their protein levels under the salt conditions. By enhancing ATP consumption in the mevalonate pathway, it is thus possible to provide an overall strong thermodynamic driving force for terpene synthesis under physiological conditions. In other studies, increasing ATP consumption has been successfully applied to drive thermodynamically unfavorable reactions (37, 38). It is also worth mentioning that although respiration rates are higher in cells grown with sodium, the intracellular ATP levels are maintained at similar levels under both salt conditions (Fig. 4A), showing a tightly regulated ATP homeostasis by balancing ATP production and consumption.

In order to accommodate the higher respiration needs, *Thraustochytrium* sp. cells seem to achieve this by shifting from carbohydrate metabolism to lipid oxidation to provide abundant reducing equivalents for the electron transport chain. Several key enzymes in the EMP pathway and pyruvate oxidation showed decreased levels in cells grown with sodium (Fig. 2). On the other hand, acyl-CoA dehydrogenase, the key enzyme in the β-oxidation pathway, had significantly higher abundance in cells grown with sodium. β-oxidation bypasses the TCA cycle to generate reducing equivalents. Not only will it provide acetyl-CoA for terpene biosynthesis, it also provides NADH and FADH_2_ to directly participate in the electron transport chain for ATP production. For every three acetyl-CoA that enter the mevalonate pathway, cells need to invest 3 ATP and 2 NADPH, which could explain why increased ATP production favors squalene synthesis in *Thraustochytrium* sp.. In plants, extracellular ATP can also contribute to the activation of the MVA pathway by phosphorylating mevalonate kinase through an ATP receptor P2K1 (39). However, it is not known whether the same mechanism exists in thraustochytrids.

It is well known that many terpenoids play important roles in stress response and osmotic adjustment (40, 41). *Thraustochytrium* sp. employs a sodium-dependent mechanism to induce higher respiration, resulting in increased terpenoids biosynthesis. However, it is currently unknown how sodium induces higher respiration. A likely mechanism could implicate the Ca^2+^-dependent energy activation mechanism. Calcium ions are important second messengers of cell metabolism, and can stimulate ATP synthesis in mitochondria by activating diverse enzymes in the electron transport chain (42). The influx of Ca^2+^ into mitochondria through the Na^+^/Ca^2+^ exchange channel in the mitochondria membrane has been shown to activate ATP generation in mouse heart cells (43). In thraustochytrids, ion exchange channels are a major response mechanism for osmotic stress (44), which could have altered intracellular Ca^2+^ levels and led to increased respiration in the mitochondria. However, the exact mechanism remains to be illustrated.

Enhancing terpene flux is critical for high terpene yield in metabolic engineering. Besides kinetic improvement, increasing overall thermodynamic driving force by increased acetyl-CoA input has led to enhanced terpene flux and high terpene yield (27, 45). We further show in this study that the mevalonate-derived terpene biosynthesis is constrained by local thermodynamics, which could be relieved by enhanced ATP investment. This strategy provides an important alternative solution for future terpene engineering efforts.

## MATERIALS AND METHODS

### Strain and medium

The *Thraustochytrium* sp. ATCC26185 strain was purchased from American Type Culture Collection (Manassas, VA, USA). Cells were cultured at 28 °C at 170 rpm in the enriched medium (EM) consisting of 30 g/L glucose, 2.5 g/L yeast extract, 2 g/L monosodium glutamate, 1 g/L KCl, 5 g/L MgSO_4_.7H_2_O, 0.1 g/L NaHCO_3_, 0.3 g/L CaCl_2_, 0.3 g/L KH_2_PO_4_, 3 mg/L FeCl_3_, 0.6 mg/L ZnSO_4_.7H_2_O, 8.6 mg/L MnSO_4_.7H_2_O, 0.26 mg/L CoCl.6H_2_O, and 0.02 mg/L CuSO_4_.5H_2_O. The seed culture was prepared by inoculating a single colony in 100 mL of EM with 25 g/L NaCl, and was grown for 2 days at 28 °C with 170 rpm. For all salt induction experiments, 0.3 OD_660nm_ of the seed culture was inoculated into 100 mL of EM with or without supplementing 5 g/L NaCl. Cell growth, squalene and total lipid contents, ATP levels, and oxygen consumption rates were measured by collecting cells at 24, 48, 60, and 72 hours.

### Squalene quantification

Two mL of cells were collected at each time point by centrifuging at 3,000 × g for 5 minutes. The cell pellet was resuspended by vortexing in 1 mL of chloroform: methanol (1:2 v/v) for 3 min. 30 μg of cedrene was added to the mixture to serve as the internal standard, followed by the addition of 1 mL acetonitrile and 2 mL hexane. The solvent mixture was vortexed for 1 minute, and centrifuged at 3,000 g for 2 min. The extracted squalene in the top hexane phase was then transferred into a new tube, and adjusted to a final volume of 2 mL using hexane. Squalene contents were analyzed by gas chromatography-mass spectrometry (GC-MS) in a Thermo Trace 1300 ISQ QD connected with a Zebron GC column (ZB-5MSplus 30 m × 0.25 mm × 0.25 μm). The GC oven temperature was held at 90 °C for 1 min, followed by a temperature increase to 300 °C at the rate of 25 °C /min, and a final temperature hold at 300 °C for 5 min. Squalene concentration was determined by calculating against a standard curve with 15 ppm of cedrene as the internal standard.

### Determination of total fatty acids

One mL of cells was centrifuged at 10000 × g for 2 minutes to collect the cell pellet. 200 μL of chloroform: methanol (2:1; v/v) and 300 μL of 0.6 M HCl-methanol solution were added into the cell pellet, followed by the addition of 30 μg of cedrene as the internal standard. The solvent mixture was vortexed for 1 min, and incubated in a 65 °C water bath for 2 hours. Following incubation, the mixture was cooled to room temperature. One mL of hexane was then added to extract the total lipids by thoroughly vortexing for 1 min. Finally, phase separation was achieved by centrifuging at 5000 rpm for 2 min. The top hexane phase was collected for GC-MS analysis in the Thermo Trace 1300 ISD QD. The initial oven temperature was set to 50 °C and maintained for 0.5 min, followed by a temperature increase to 180 °C at the rate of 25 °C/min and hold for 5 min. The temperature was raised again to 250°C at the rate of 5 °C/min and held at 250 °C for 10 min. The amount of fatty acid components (C14, C15, C16, C17, C18, and C22) were calculated against the internal standard cedrene in the GC-MS analysis. Total fatty acids were calculated by the sum of all lipid components.

### Shotgun proteomics

Comparative proteomics was conducted by collecting *Thraustochytrium* sp. cells grown with or without 5 g/L of NaCl in the EM. About 10 OD of cells were collected at the time point of 60 hours by centrifuging at 10,000 × g for 2 min. The proteomic sample preparation and mass spectrometry analysis in a Thermo LTQ Orbitrap XL mass spectrometer were conducted following the same protocol in our previous study (46). The raw data collected from MS/MS analysis was searched and analyzed using programs integrated in the PatternLab for Proteomics (47). The peptide information was obtained by searching against the database obtained from translating our transcriptomic data (not published). The proteomics data has been deposited to the ProteomeXchange Consortium with the dataset identifier PXD021931 (48).

### *in-silico* analysis of pathway thermodynamics

The thermodynamic feasibility of squalene synthesis from both glycolysis and the β-oxidation of a short chain fatty acid (hexanoyl-CoA) was evaluated using the pathway analysis tool PathParser (28). PathParser seeks to solve a max-min problem by maximizing the Gibbs energy change (ΔG’) of the most thermodynamically unfavorable reaction in the pathway, which can be formulated as:

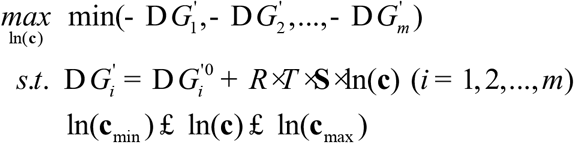

where m is the number of reactions in a pathway, **c** is the concentration vector of metabolites involved, and **S** is the stoichiometric matrix of the pathway reactions. The standard Gibbs free energies (ΔG’^m^) were searched in eQuilibrator database (34) with intracellular pH and ionic strength set to 7.5 and 0.2 M, respectively. The lower and upper bounds of metabolite concentrations were constrained to range from 1 μM to 10 mM, following the approximate concentration scale of cellular metabolites (49). In addition, cofactor concentrations were fixed to approximate levels found in eukaryotic cells (30), which allowed a proper evaluation of target pathway thermodynamics with encoded constraints from the wider metabolic network (29). Specifically, the following cofactor concentrations were defined: ATP = 0.005 M, ADP = 0.0005 M, NAD = 0.001 M, NADH = 0.0001 M, NADPH = 0.0001 M, NADP = 0.0001 M, orthophosphate = 0.01 M, Coenzyme A: 0.001 M, CO_2_: 0.00001 M, and diphosphate: 0.001 M.

### ATP determination

ATP levels were monitored throughout the cell growth. About 1 OD of cells were collected by centrifuging at maximum speed for 15 s at 4 °C. Cells were quenched by adding 1 mL of ice-cold methanol, followed by incubation for 1 hour at −30 °C. The extracted ATP in the methanol solution was then dried in a speed-vacuum, and dissolved in deionized water. ATP levels were determined using the ATP Determination Kit (Molecular Probes) according to the manufacturer’s instruction.

### Oxygen consumption rate

Cell respiration rates were determined by measuring oxygen consumption rates in a Clark type oxygen electrode (Oxygraph System,

Hansatech Instruments). Two mL of cells grown with or without NaCl were collected at the time points of 24, 48, 60, and 72 hours. No oxygen consumption was observed in the blank EM with or without the addition of NaCl. All cells were diluted to a density of ~ 0.5 OD and monitored for 2 min at room temperature for oxygen consumption rates. The final oxygen consumption rates were normalized by cell optical density.

## ACKNOWLEDGMENTS

This work was supported by Miami University start-up fund to X.W. and the scholarship from Chinese Scholarship Council to A.Z.. We would like to thank Dr. Andor Kiss and Ms. Xiaoyun Deng of the Center for Bioinformatics and Functional Genomics at Miami University for instrument support. This work was authored in part by the National Renewable Energy Laboratory, operated by Alliance for Sustainable Energy, LLC, for the U.S. Department of Energy (DOE) under Contract No. DE-AC36-08GO28308. Funding provided by Laboratory Directed Research and Development program. The views expressed in the article do not necessarily represent the views of the DOE or the U.S. Government. The U.S. Government retains and the publisher, by accepting the article for publication, acknowledges that the U.S. Government retains a nonexclusive, paid-up, irrevocable, worldwide license to publish or reproduce the published form of this work, or allow others to do so, for U.S. Government purposes.

