## Supplemental information for "ATP drives efficient terpene biosynthesis in marine thraustochytrids"

**Table S1 Fatty acid production of *Thraustochytrium* sp. cultured under different NaCl conditions.**

| | | **C14:0 Myristic acid** | | **C15:0 Pentadecylic acid** | | **C16:0 Palmitic acid** | | **C17:0 Margaric acid** | | **C18:0 Stearic acid** | | **C22:6 Docosahexaenoic acid** | |
|---|---|---|---|---|---|---|---|---|---|---|---|---|---|
| **Time (hour)** | Group | Titer (mg/L) | Specific productivity (mg/L/day/OD) | Titer (mg/L) | Specific productivity (mg/L/day/OD) | Titer (mg/L) | Specific productivity (mg/L/day/OD) | Titer (mg/L) | Specific productivity (mg/L/day/OD) | Titer (mg/L) | Specific productivity (mg/L/day/OD) | Titer (mg/L) | Specific productivity (mg/L/day/OD) |
| **24** | NaCl-0 | 0.00 | 0.00 | 15.98 | 5.97 | 87.18 | 32.70 | 2.98 | 1.12 | 26.78 | 10.06 | 36.94 | 13.83 |
| | NaCl-5 | 0.00 | 0.00 | 8.61 | 2.35 | 62.84 | 17.07 | 1.84 | 0.50 | 17.39 | 4.71 | 34.47 | 9.37 |
| **48** | NaCl-0 | 16.39 | 1.04 | 37.25 | 2.35 | 138.42 | 8.77 | 5.88 | 0.37 | 29.27 | 1.86 | 57.91 | 3.66 |
| | NaCl-5 | 21.12 | 1.40 | 52.57 | 3.49 | 137.19 | 9.11 | 6.15 | 0.41 | 30.11 | 2.00 | 66.31 | 4.41 |
| **60** | NaCl-0 | 21.95 | 0.70 | 50.71 | 1.62 | 153.69 | 4.88 | 8.14 | 0.26 | 32.34 | 1.01 | 72.69 | 2.31 |
| | NaCl-5 | 27.61 | 0.95 | 87.15 | 3.00 | 149.21 | 5.14 | 12.27 | 0.42 | 32.75 | 1.13 | 92.00 | 3.17 |
| **72** | NaCl-0 | 27.57 | 0.84 | 58.42 | 1.78 | 164.08 | 4.99 | 9.24 | 0.28 | 39.23 | 1.19 | 86.61 | 2.63 |
| | NaCl-5 | 27.31 | 0.88 | 83.68 | 2.71 | 136.63 | 4.43 | 12.49 | 0.40 | 35.75 | 1.16 | 94.76 | 3.07 |

**Table S2 Thermodynamic analysis of squalene synthesis from glucose**. Standard Gibbs free energies ($\Delta G'^m$) were obtained from eQuilibrator database (1). Optimal $\Delta G'$ numbers were calculated from the PathParser (2). The cofactor coenzyme A was fixed to 1 mM in the analysis.

| **Enzyme** | **Enzyme ID** | **Reaction** | **ΔG'm (kJ mol-1)** | **Optimal ΔG' (kJ mol-1)** | **Occurrence** |
|---|---|---|---|---|---|
| Glucokinase | HXK | D-Glucose + ATP = D-Glucose 6-phosphate + ADP | -19.4 | -15.6 | 9 |
| Glucose-6-phosphate isomerase | PGI | D-Glucose 6-phosphate = D-Fructose 6-phosphate | 2.5 | -6.6 | 9 |
| 6-phosphofructokinase | PFK | ATP + D-Fructose 6-phosphate = ADP + Fructose-1,6-bisphosphate | -16.6 | -10.0 | 9 |
| Fructose-bisphosphate aldolase | FBA | Fructose-1,6-bisphosphate = Glycerone phosphate + D-Glyceraldehyde 3-phosphate | 2.2 | -3.6 | 9 |
| Triose-phosphate isomerase | TPI | Glycerone phosphate = D-Glyceraldehyde 3-phosphate | 5.5 | -2.9 | 9 |
| GAP dehydrogenase | GAPDH | Pi + NAD+ + D-Glyceraldehyde 3-phosphate = NADH + 1,3-Bisphosphoglycerate | 39.4 | -3.2 | 18 |
| Phosphoglycerate kinase | PGK | ADP + 1,3-Bisphosphoglycerate = ATP + D-Glycerate-3-phosphate | -37.4 | -5.3 | 18 |
| Phosphoglycerate mutase | PGM | D-Glycerate-3-phosphate = D-Glycerate-2-phosphate | 8.2 | -5.5 | 18 |
| Enolase | ENO | D-Glycerate-2-phosphate = Phosphoenolpyruvate + H2O | -8.2 | -8.7 | 18 |
| Pyruvate kinase | PK | ADP + Phosphoenolpyruvate = Pyruvate + ATP | -49.6 | -30.3 | 18 |
| Pyruvate dehydrogenase | PDH | NAD+ + CoA + Pyruvate = NADH + $CO_2$ + Acetyl-CoA + H+ | -71 | -76.8 | 18 |
| Acetyl-CoA acyltransferase | ACAT | 2 Acetyl-CoA = CoA + Acetoacetyl-CoA | 17.3 | -1.7 | 6 |
| Hydroxymethylglutaryl-CoA synthase | HMGS | H2O + Acetyl-CoA + Acetoacetyl-CoA = CoA + (S)-3-Hydroxy-3-methylglutaryl-CoA | -14.2 | -11.0 | 6 |
| Hydroxymethylglutaryl-CoA reductase | HMGR | 2 NADPH + 2 H+ + (S)-3-Hydroxy-3-methylglutaryl-CoA = 2 NADP+ + CoA + (R)-Mevalonate | -10.8 | -10.8 | 6 |
| Mevalonate kinase | MVK | ATP + (R)-Mevalonate = ADP + (R)-5-Phosphomevalonate | -10.7 | -12.6 | 6 |
| phosphomevalonate kinase | PMK | ATP + (R)-5-Phosphomevalonate = ADP + (R)-5-Diphosphomevalonate | -1.3 | -9.2 | 6 |
| mevalonate diphosphate decarboxylase | MDD | ATP + (R)-5-Diphosphomevalonate = ADP + Orthophosphate + $CO_2$ + Isopentenyl diphosphate | -43.4 | -44.6 | 6 |
| IPP isomerase | IDI | Isopentenyl diphosphate = Dimethylallyl diphosphate | -1.1 | -3.3 | 2 |
| FPP synthase | FPS | 2 Isopentenyl diphosphate + Dimethylallyl diphosphate = 2 Diphosphate + Farnesyl diphosphate | -33.4 | -32.3 | 2 |
| squalene synthase | SQS | NADPH + H+ + 2 Farnesyl diphosphate = NADP+ + 2 Diphosphate + Squalene | -20 | -19.4 | 1 |

**Table S3 Thermodynamic analysis of squalene synthesis from the short-chain fatty acid hexanoyl-CoA**. Standard Gibbs free energies ($\Delta G'^{m}$) were obtained from eQuilibrator database (1). Optimal $\Delta G'$ numbers were calculated from the PathParser (2). The cofactor coenzyme A was fixed to 1 mM in the analysis.

| **Enzyme** | **Enzyme ID** | **Reaction** | **$\Delta G'^{m}$ (kJ mol$^{-1}$)** | **Optimal $\Delta G'$ (kJ mol$^{-1}$)** | **Occurrence** |
|---|---|---|---|---|---|
| acyl-CoA dehydrogenase | ACAD | Hexanoyl-CoA = trans-Hex-2-enoyl-CoA + 2 e- | -8 | -4.1 | 6 |
| enoyl-CoA hydratase | ECH | H2O + trans-Hex-2-enoyl-CoA = (S)-Hydroxyhexanoyl-CoA | -1 | -1.61 | 6 |
| β-hydroxyacyl-CoA dehydrogenase | HADH | NAD+ + (S)-Hydroxyhexanoyl-CoA = NADH + H+ + 3-Oxohexanoyl-CoA | 21 | -1.11 | 6 |
| acyl-CoA acetyltransferase | KAT | CoA + 3-Oxohexanoyl-CoA = Acetyl-CoA + Butanoyl-CoA | -34.9 | -18.6 | 6 |
| acyl-CoA dehydrogenase | ACAD | Butanoyl-CoA = Crotonoyl-CoA + 2 e- | -13.2 | -10.86 | 6 |
| enoyl-CoA hydratase | ECH | H2O + Crotonoyl-CoA = (S)-3-Hydroxybutanoyl-CoA | -3.8 | -4.96 | 6 |
| β-hydroxyacyl-CoA dehydrogenase | HADH | NAD+ + (S)-3-Hydroxybutanoyl-CoA = NADH + H+ + Acetoacetyl-CoA | 14.6 | -3.6 | 6 |
| acyl-CoA acetyltransferase | KAT | CoA + Acetoacetyl-CoA = 2 Acetyl-CoA | -26 | 1.52E-12 | 6 |
| Acetyl-CoA acyltransferase | ACAT | 2 Acetyl-CoA = CoA + Acetoacetyl-CoA | 26 | -1.52E-12 | 6 |
| Hydroxymethylglutaryl-CoA synthase | HMGS | H2O + Acetyl-CoA + Acetoacetyl-CoA = CoA + (S)-3-Hydroxy-3-methylglutaryl-CoA | -21.3 | -17.2 | 6 |
| Hydroxymethylglutaryl-CoA reductase | HMGR | 2 NADPH + 2 H+ + (S)-3-Hydroxy-3-methylglutaryl-CoA = 2 NADP+ + CoA + (R)-Mevalonate | -16.2 | -16.2 | 6 |
| Mevalonate kinase | MVK | ATP + (R)-Mevalonate = ADP + (R)-5-Phosphomevalonate | -16.1 | -19.0 | 6 |
| phosphomevalonate kinase | PMK | ATP + (R)-5-Phosphomevalonate = ADP + (R)-5-Diphosphomevalonate | -1.9 | -14.7 | 6 |
| mevalonate diphosphate decarboxylase | MDD | ATP + (R)-5-Diphosphomevalonate = ADP + Orthophosphate + $CO_2$ + Isopentenyl diphosphate | -65.1 | -64.9 | 6 |
| IPP isomerase | IDI | Isopentenyl diphosphate = Dimethylallyl diphosphate | -1.6 | -5.0 | 2 |
| FPP synthase | FPS | 2 Isopentenyl diphosphate + Dimethylallyl diphosphate = 2 Diphosphate + Farnesyl diphosphate | -50.2 | -49.5 | 2 |
| squalene synthase | SQS | NADPH + H+ + 2 Farnesyl diphosphate = NADP+ + 2 Diphosphate + Squalene | -30 | -29.0 | 1 |

**Table S4 Optimal metabolite concentrations for the maximum of thermodynamic driving force for squalene synthesis.** The metabolite concentration range is between 0.001 mM and 10 mM. The cofactor coenzyme A was fixed to 1 mM in the analysis.

| Glucose to Squalene | | Hexanoyl-CoA to Squalene | |
|---|---|---|---|
| **Metabolite** | **Optimized concentration (mM)** | **Metabolite** | **Optimized concentration (mM)** |
| Glucose | 0.0164 | Hexanoyl-CoA | 0.392 |
| ATP | 5 | transHex2enoyl-CoA | 1.924 |
| G6P | 0.770 | Hydroxyhexanoyl-CoA | 1.490 |
| ADP | 0.5 | NAD | 1 |
| F6P | 0.020 | NADH | 0.1 |
| FBP | 2.800 | Oxohexanoyl-CoA | 0.00198 |
| GAP | 0.0958 | CoA | 1 |
| DHAP | 2.868 | Acetyl-CoA | 8.059 |
| Pi | 10 | Butanoyl-CoA | 0.179 |
| NAD | 1 | Crotonoyl-CoA | 0.463 |
| 1,3BPG | 0.00179 | Hydroxybutanoyl-CoA | 0.293 |
| NADH | 0.1 | Acetoacetyl-CoA | 0.00181 |
| 3PG | 0.115 | Hydroxymethylglutaryl-CoA | 0.0774 |
| 2PG | 0.00732 | NADPH | 0.1 |
| PEP | 0.00664 | NADP | 0.1 |
| Pyr | 0.0324 | Mevalonate | 0.0771 |
| CoA | 1 | ATP | 5 |
| $CO_2$ | 0.01 | ADP | 0.5 |
| Acetyl-CoA | 9.99 | Phosphomevalonate | 0.241 |
| Acetoacetyl-CoA | 0.001 | Diphosphomevalonate | 0.0136 |
| Hydroxymethylglutaryl-CoA | 0.0707 | Pi | 10 |
| NADPH | 0.1 | $CO_2$ | 0.01 |
| NADP | 0.1 | Isopentenyl PP | 1.463 |
| Mevalonate | 0.07 | Dimethylallyl PP | 0.0262 |
| Phosphomevalonate | 0.232 | PPi | 1 |
| Diphosphomevalonate | 0.0189 | Farnesyl PP | 0.128 |
| Isopentenyl PP | 0.920 | Squalene | 0.172 |
| Dimethylallyl PP | 0.0175 | | |
| PPi | 1 | | |
| Farnesyl PP | 0.127 | | |
| Squalene | 0.129 | | |

**Table S5 Thermodynamic analysis of squalene synthesis from glucose**. Standard Gibbs free energies ($\Delta G'^{m}$) were obtained from eQuilibrator database (1). Optimal $\Delta G'$ numbers were calculated from the PathParser (2). The cofactor coenzyme A was fixed to 5 mM in the analysis.

| Enzyme | Enzyme ID | Reaction | $\Delta G'^{m}$ (kJ $mol^{-1}$) | Optimal $\Delta G'$ (kJ $mol^{-1}$) | Occurrence |
|---|---|---|---|---|---|
| Glucokinase | HXK | D-Glucose + ATP = D-Glucose 6-phosphate + ADP | -19.4 | -16.6 | 9 |
| Glucose-6-phosphate isomerase | PGI | D-Glucose 6-phosphate = D-Fructose 6-phosphate | 2.5 | -5.1 | 9 |
| 6-phosphofructokinase | PFK | ATP + D-Fructose 6-phosphate = ADP + Fructose-1,6-bisphosphate | -16.6 | -12.4 | 9 |
| Fructose-bisphosphate aldolase | FBA | Fructose-1,6-bisphosphate = Glycerone phosphate + D-Glyceraldehyde 3-phosphate | 2.2 | -2.4 | 9 |
| Triose-phosphate isomerase | TPI | Glycerone phosphate = D-Glyceraldehyde 3-phosphate | 5.5 | -1.1 | 9 |
| GAP dehydrogenase | GAPDH | Pi + NAD+ + D-Glyceraldehyde 3-phosphate = NADH + 1,3-Bisphosphoglycerate | 39.4 | -1.5 | 18 |
| Phosphoglycerate kinase | PGK | ADP + 1,3-Bisphosphoglycerate = ATP + D-Glycerate-3-phosphate | -37.4 | -5.1 | 18 |
| Phosphoglycerate mutase | PGM | D-Glycerate-3-phosphate = D-Glycerate-2-phosphate | 8.2 | -4.4 | 18 |
| Enolase | ENO | D-Glycerate-2-phosphate = Phosphoenolpyruvate + H2O | -8.2 | -8.3 | 18 |
| Pyruvate kinase | PK | ADP + Phosphoenolpyruvate = Pyruvate + ATP | -49.6 | -34.6 | 18 |
| Pyruvate dehydrogenase | PDH | NAD+ + CoA + Pyruvate = NADH + CO2 + Acetyl-CoA + H+ | -71 | -84.9 | 18 |
| Acetyl-CoA acyltransferase | ACAT | 2 Acetyl-CoA = CoA + Acetoacetyl-CoA | 17.3 | 1 | 6 |
| Hydroxymethylglutaryl-CoA synthase | HMGS | H2O + Acetyl-CoA + Acetoacetyl-CoA = CoA + (S)-3-Hydroxy-3-methylglutaryl-CoA | -14.2 | -8.6 | 6 |
| Hydroxymethylglutaryl-CoA reductase | HMGR | 2 NADPH + 2 H+ + (S)-3-Hydroxy-3-methylglutaryl-CoA = 2 NADP+ + CoA + (R)-Mevalonate | -10.8 | -8.3 | 6 |
| Mevalonate kinase | MVK | ATP + (R)-Mevalonate = ADP + (R)-5-Phosphomevalonate | -10.7 | -12.3 | 6 |
| phosphomevalonate kinase | PMK | ATP + (R)-5-Phosphomevalonate = ADP + (R)-5-Diphosphomevalonate | -1.3 | -8.1 | 6 |
| mevalonate diphosphate decarboxylase | MDD | ATP + (R)-5-Diphosphomevalonate = ADP + Orthophosphate + CO2 + Isopentenyl diphosphate | -43.4 | -47.2 | 6 |
| IPP isomerase | IDI | Isopentenyl diphosphate = Dimethylallyl diphosphate | -1.1 | -2.2 | 2 |
| FPP synthase | FPS | 2 Isopentenyl diphosphate + Dimethylallyl diphosphate = 2 Diphosphate + Farnesyl diphosphate | -33.4 | -31.8 | 2 |
| squalene synthase | SQS | NADPH + H+ + 2 Farnesyl diphosphate = NADP+ + 2 Diphosphate + Squalene | -20 | -19.3 | 1 |

**Table S6 Thermodynamic analysis of squalene synthesis from the short-chain fatty acid hexanoyl-CoA**. Standard Gibbs free energies ($\Delta G'^m$) were obtained from eQuilibrator database (1). Optimal $\Delta G'$ numbers were calculated from the PathParser (2). The cofactor coenzyme A was fixed to 5 mM in the analysis.

| Enzyme | Enzyme ID | Reaction | ΔG'm (kJ mol-1) | Optimal ΔG' (kJ mol-1) | Occurrence |
|---|---|---|---|---|---|
| acyl-CoA dehydrogenase | ACAD | Hexanoyl-CoA = trans-Hex-2-enoyl-CoA + 2 e- | -8 | -4.3 | 6 |
| enoyl-CoA hydratase | ECH | H2O + trans-Hex-2-enoyl-CoA = (S)-Hydroxyhexanoyl-CoA | -1 | -1.2 | 6 |
| β-hydroxyacyl-CoA dehydrogenase | HADH | NAD+ + (S)-Hydroxyhexanoyl-CoA = NADH + H+ + 3-Oxohexanoyl-CoA | 21 | -0.3 | 6 |
| acyl-CoA acetyltransferase | KAT | CoA + 3-Oxohexanoyl-CoA = Acetyl-CoA + Butanoyl-CoA | -34.9 | -22.1 | 6 |
| acyl-CoA dehydrogenase | ACAD | Butanoyl-CoA = Crotonoyl-CoA + 2 e- | -13.2 | -11.6 | 6 |
| enoyl-CoA hydratase | ECH | H2O + Crotonoyl-CoA = (S)-3-Hydroxybutanoyl-CoA | -3.8 | -5.5 | 6 |
| β-hydroxyacyl-CoA dehydrogenase | HADH | NAD+ + (S)-3-Hydroxybutanoyl-CoA = NADH + H+ + Acetoacetyl-CoA | 14.6 | -4.2 | 6 |
| acyl-CoA acetyltransferase | KAT | CoA + Acetoacetyl-CoA = 2 Acetyl-CoA | -26 | -1.5 | 6 |
| Acetyl-CoA acyltransferase | ACAT | 2 Acetyl-CoA = CoA + Acetoacetyl-CoA | 26 | 1. 5 | 6 |
| Hydroxymethylglutaryl-CoA synthase | HMGS | H2O + Acetyl-CoA + Acetoacetyl-CoA = CoA + (S)-3-Hydroxy-3-methylglutaryl-CoA | -21.3 | -12.2 | 6 |
| Hydroxymethylglutaryl-CoA reductase | HMGR | 2 NADPH + 2 H+ + (S)-3-Hydroxy-3-methylglutaryl-CoA = 2 NADP+ + CoA + (R)-Mevalonate | -16.2 | -13 | 6 |
| Mevalonate kinase | MVK | ATP + (R)-Mevalonate = ADP + (R)-5-Phosphomevalonate | -16.1 | -18.7 | 6 |
| phosphomevalonate kinase | PMK | ATP + (R)-5-Phosphomevalonate = ADP + (R)-5-Diphosphomevalonate | -1.9 | -13.4 | 6 |
| mevalonate diphosphate decarboxylase | MDD | ATP + (R)-5-Diphosphomevalonate = ADP + Orthophosphate + $CO_2$ + Isopentenyl diphosphate | -65.1 | -68.2 | 6 |
| IPP isomerase | IDI | Isopentenyl diphosphate = Dimethylallyl diphosphate | -1.6 | -4.1 | 2 |
| FPP synthase | FPS | 2 Isopentenyl diphosphate + Dimethylallyl diphosphate = 2 Diphosphate + Farnesyl diphosphate | -50.2 | -48 | 2 |
| squalene synthase | SQS | NADPH + H+ + 2 Farnesyl diphosphate = NADP+ + 2 Diphosphate + Squalene | -30 | -29 | 1 |

**Table S7 Optimal metabolite concentrations for the maximum of thermodynamic driving force for squalene synthesis.** The metabolite concentration range is between 0.001 mM and 10 mM. The cofactor coenzyme A was fixed to 5 mM in the analysis.

| Glucose to Squalene | | Hexanoyl-CoA to Squalene | |
|---|---|---|---|
| **Metabolite** | **Optimized concentration (mM)** | **Metabolite** | **Optimized concentration (mM)** |
| Glucose | 0.0155 | Hexanoyl-CoA | 0.316 |
| ATP | 5 | transHex2enoyl-CoA | 1.385 |
| G6P | 0.482 | Hydroxyhexanoyl-CoA | 1.286 |
| ADP | 0.5 | NAD | 1 |
| F6P | 0.0224 | NADH | 0.1 |
| FBP | 1.242 | Oxohexanoyl-CoA | 0.00236 |
| GAP | 0.118 | CoA | 5 |
| DHAP | 1.661 | Acetyl-CoA | 9.99 |
| Pi | 10 | Butanoyl-CoA | 0.204 |
| NAD | 1 | Crotonoyl-CoA | 0.385 |
| 1,3BPG | 0.00308 | Hydroxybutanoyl-CoA | 0.197 |
| NADH | 0.1 | Acetoacetyl-CoA | 0.001 |
| 3PG | 0.207 | Hydroxymethylglutaryl-CoA | 0.0784 |
| 2PG | 0.0164 | NADPH | 0.1 |
| PEP | 0.016 | NADP | 0.1 |
| Pyr | 0.033 | Mevalonate | 0.0579 |
| CoA | 5 | ATP | 5 |
| $CO_2$ | 0.01 | ADP | 0.5 |
| Acetyl-CoA | 10 | Phosphomevalonate | 0.2012 |
| Acetoacetyl-CoA | 0.001 | Diphosphomevalonate | 0.0197 |
| Hydroxymethylglutaryl-CoA | 0.0584 | Pi | 10 |
| NADPH | 0.1 | $CO_2$ | 0.01 |
| NADP | 0.1 | Isopentenyl PP | 0.555 |
| Mevalonate | 0.0534 | Dimethylallyl PP | 0.0282 |
| Phosphomevalonate | 0.213 | PPi | 1 |
| Diphosphomevalonate | 0.0333 | Farnesyl PP | 0.114 |
| Isopentenyl PP | 0.336 | Squalene | 0.141 |
| Dimethylallyl PP | 0.0477 | | |
| PPi | 1 | | |
| Farnesyl PP | 0.114 | | |
| Squalene | 0.124 | | |
